# Disruption of *AtHAK/KT/KUP9* enhances plant cesium accumulation under low potassium supply

**DOI:** 10.1101/2020.02.05.931527

**Authors:** Laure Genies, Ludovic Martin, Satomi Kanno, Serge Chiarenza, Loïc Carasco, Virginie Camilleri, Alain Vavasseur, Pascale Henner, Nathalie Leonhardt

## Abstract

Understanding molecular mechanisms which underlie transport of cesium (Cs^+^) in plants is important to limit entry of its radioisotopes from contaminated area to the food chain. The potentially toxic element Cs^+^, which is not involved in any biological process, is chemically closed to the macronutrient potassium (K^+^). Among the multiple K^+^ carriers, the high-affinity K^+^ transporters family HAK/KT/KUP is thought to be relevant in mediating opportunistic Cs^+^ transport. On the 13 *KUP* identified in *Arabidopsis thaliana*, only *HAK5*, the major contributor to root K^+^ acquisition under low K^+^ supply, has been functionally demonstrated to be involved in Cs^+^ uptake *in planta*. In the present study, we showed that accumulation of Cs^+^ increased by up to 30% in two *A. thaliana* mutant lines lacking *KUP9* and grown under low K^+^ supply. Since further experiments revealed that Cs^+^ release from contaminated plants to the external medium is proportionally lower in the two *kup9* mutants, we proposed that *KUP9* disruption could impair Cs^+^ efflux. By contrast, we did not measure significant impairment of K^+^ status in *kup9* mutants suggesting that *KUP9* disruption does not alter substantially K^+^ transport in experimental conditions used here. Putative primary role of KUP9 in plants is further discussed.

## INTRODUCTION

Cesium is an alkali metal occurring generally at low concentration in soil solution where it is present predominantly as the monovalent cation Cs^+^ (Greenwood & Earnshaw, 1984). Although it has not been involved in any biological process to date, Cs^+^ is taken up by plants and can even be toxic (Hampton et al., 2004). Due to its low environmental concentration, the chemical toxicity of stable Cs^+^ is rarely relevant in natural conditions (White & Broadley, 2000). By contrast, the radiological threat of ^134^Cs and ^137^Cs, two radioisotopes of Cs^+^ originated from nuclear activities, is a concern for environment and human health in contaminated areas. These two β- and γ-emitting radionuclides were among those unintentionally released at harmful levels during nuclear accidents including those at Fukushima (Japan, 2011) and Chernobyl (Ukraine, 1986) (Steinhauser, Brandl & Johnson, 2014) and remained monitored due to their relative long half-lives (2.06 and 30.17 years for ^134^Cs and ^137^Cs respectively). Contaminated food is regarded as a major source of radionuclides exposure for humans after the initial phase of a nuclear accident (Hamada & Ogino, 2012). Therefore, understanding transfer of radiocesium in plants remains a challenging question for reducing its transfer into the food chain through development of “safe crops” which accumulate less amounts of radionuclides, or through phytoremediation strategies to remove Cs^+^ from soils (White & Broadley, 2000).

To date, it is commonly admitted that plants do not discriminate significantly stable and radioactive isotopes of Cs^+^ (White & Broadley, 2000) which are used indifferently for uptake experiments. Cs^+^ shares closed chemical properties with the macronutrient K^+^ (Bowen, 1979) and early studies demonstrated that mechanisms underlying Cs^+^ and K^+^ uptake in plants were also similar (Collander, 1941; Epstein & Hagen, 1952). As for K^+^, Cs^+^ uptake in plants can be divided into two major systems: a high-affinity transport system (HATS) operating for external Cs^+^ concentration in the micromolar range and a low-affinity transport system (LATS) operating in the millimolar range (Bange & Overstreet, 1960; Shaw & Bell, 1989). In addition, several studies on different plant species described a competitive effect of external K^+^ on Cs^+^ uptake (Kondo et al., 2015; Middleton, Handley & Overstreet, 1960; Sacchi, Espen, Nocito & Cocucci, 1997; Smolders, Kiebooms, Buysse & Merckx, 1996). Since Cs^+^ has no relevant physiological role in plants and given the above-mentioned evidences suggesting that K^+^ and Cs^+^ share the same transport mechanisms, it is generally assumed that Cs^+^ entry in plants is mainly mediated through K^+^ transporters (White & Broadley, 2000).

Multiple transport systems are involved in plant K^+^ acquisition and contribution of each individual system depends on the external K^+^ concentration (Alemán, Nieves-Cordones, Martínez & Rubio, 2011). In the same way, K^+^ supply also affects contribution of each pathway mediating Cs^+^ uptake. In the model plant *Arabidopsis thaliana*, Non-selective Cation Channels (NSCC) are assumed to mediate a significant part of Cs^+^ uptake under sufficient K^+^-supply (Hampton, Broadley & White, 2005; White & Broadley, 2000). The *Arabidopsis* Zinc-Induced-Facilitator-Like-2 (ZIFL2) carrier is also involved in Cs^+^ partitioning in K^+^-replete plants (Remy et al., 2015). In contrast, the Shaker channel AKT1 which is a major contributor of K^+^ uptake in roots of *A. thaliana* (Hirsch, Lewis, Spalding & Sussman, 1998), is Cs^+^-sensitive (Bertl, Reid, Sentenac & Slayman, 1997) but is not relevant for plant Cs^+^ uptake (Broadley, Escobar-Gutiérrez, Bowen, Willey & White, 2001).

In K^+^-starved plants, transporters encoded by the *HAK/KT/KUP* genes family (named *KUP* in the following) have been pointed out as a relevant pathway for Cs^+^ (Rubio, Guillermo & Rodríguez-Navarro, 2000; White & Broadley, 2000). This statement is supported by several evidences among those the role of bacterial KUP transporters in Cs^+^ uptake (Bossemeyer, Schlösser & Bakker, 1989), the demonstration that Cs^+^ uptake in maize roots is mediated by K^+^ HATS (Sacchi et al., 1997) which involve members of the *KUP* family (Alemán et al., 2011), the transport of Cs^+^ through plants KUP transporter expressed in yeast (*AtHAK5*, Rubio et al., 2000) and in bacteria (*AtKUP9*, Kobayashi, Uozumi, Hisamatsu & Yamagami, 2010).

In *A. thaliana*, 13 genes encode for KUP transporters (Mäser et al., 2001) among those only the high-affinity K^+^ transporter HAK5 (Gierth, Mäser & Schroeder, 2005; Rubio et al., 2000) has been demonstrated to be functionally involved in Cs^+^ uptake (Qi et al., 2008). In the present study, we investigated the role of the KUP9 transporter in Cs^+^ accumulation in *A. thaliana*. Although this transporter has been demonstrated to be involved in Cs^+^ influx when expressed in an *Escherichia coli* mutant defective in K^+^ transport systems (Kobayashi et al., 2010), no role in plant Cs^+^ accumulation has been reported for KUP9 up to now. Using *A. thaliana* mutants disrupted in *KUP9* gene, we provide *in planta* evidence that KUP9 play a significant role in Cs^+^ accumulation whereas its disruption does not alter substantially K^+^ homeostasis.

## MATERIAL & METHODS

### Plant material

*Arabidopsis thaliana* ecotype Columbia-0 (Col-0) was used as the wild-type. Consequences of a disruption of *KUP9* was studied using two independent *kup9* T-DNA insertion lines obtained from the SAIL (N862313) and the SALK (N670022) collections respectively. T-DNA insertion was checked and homozygous lines were identified using combination of T-DNA border primers and gene specific primers as outlined in the online protocol “Screening SALK T-DNA mutagenesis lines” (University of Wisconsin, Madison Knockout Facility and Ohio State University, Arabidopsis Biological Resource Center https://www.mcdb.ucla.edu/Research/Goldberg/HC70AL_S06/pdf/Expt6protocol.pdf). In the following, the SALK_108080C mutant line with T-DNA insertion located in exon 9 and the SAIL_211_E04 mutant line with T-DNA insertion located in intron 2 are named respectively *kup9-1* (Tenorio-Berrío et al., 2018) and *kup9-3* (**Fig. 1**).

**Figure 1:**
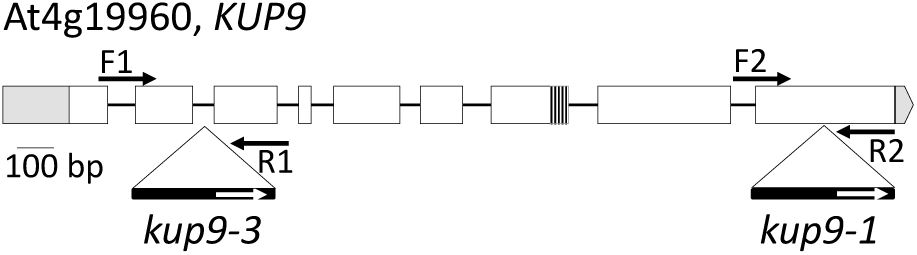
T-DNA insertion sites in *kup9-1* (SALK_108080C) and *kup9-3* (SAIL_211_E04) mutant lines. White boxes represent exons whereas dark lines between boxes denote introns of *AtKUP9* gene. Splicing variant is depicted by a grey box in exon 7. T-DNA insertions are outlined by large triangle and black arrows represent gene specific primers used for mutant lines checking (F1: GGAGATTTAGGGACGTCTCCATTGTATGTG, R1: TCCTCATCACTACGGTGCTGATTCG; F2: CCTACAGCAGCACGTATTCCGTCAAC, R2: CGGTGTTCCCCATTATATGAACAACACCTG).

### Growth conditions

*Arabidopsis* seeds were surface-sterilized using a mix of 70% ethanol (v/v) / 0.05% SDS (v/v) and rinsed in ethanol 96% before sowing in Petri dishes (120 mm * 120 mm) on a half-strength Murashige and Skoog medium (MS½, Murashige & Skoog, 1962) containing 1% (w/v) agar and 1% (w/v) sucrose. The sowing boxes were placed at 4 °C during 48 h before transfer in a growth chamber set to 23 °C, 50% relative humidity with 8 h/16 h day/night cycle. Then, 7 days-old seedlings were transferred on sand (Zolux) watered with a solution (pH 5.8) containing 1.1 mM MgSO_4_, 805 µM Ca(NO_3_)_2_, 2 mM KNO_3_, 60 µM K_2_HPO_4_, 695 µM KH_2_PO_4_ and micronutrients (3.6 µM MnSO_4_, 74 nM (NH_4_)_6_Mo_7_O_24_, 3 µM ZnSO_4_, 9.25 µM H_3_BO_3_, 785 nM CuSO_4_, 20 µM Na_2_EDTA and 20 µM FeSO_4_). Finally, 21 days-old plants were transferred on 1 L of this solution during 3-5 days for acclimation to hydroponics growing.

In subsequent experiments, basic composition of the nutritive media with controlled K^+^ and Cs^+^ content is: 0,75 mM MgSO_4_, 2 mM Ca(NO_3_)_2_, 0,5 mM H_3_PO_4_, 3,5 mM MES, 10 µM Fe-EDTA, 3,6 µM MnSO_4_, 74 nM (NH_4_)_6_Mo_7_O_24_, 3 µM ZnSO_4_, 9,25 µM H_3_BO_3_, 785 nM CuSO_4_, pH adjusted to 5.8 with NMDG (and 1% (w/v) agar for agar plates only). K^+^ and Cs^+^ concentrations on the different conditions were adjusted with KCl and CsCl.

### K^+^, Rb^+^ and Cs^+^ accumulation in seedlings

Accumulation experiments in plants were performed in hydroponics conditions with the nutritive solution described above. Protocol for long-term Cs^+^ accumulation assays in hydroponics conditions is described elsewhere (Genies et al., 2017). Briefly, after acclimation to hydroponics, 25 days-old (± 1 day) plants were grown in 10 or 3000 µM K^+^-solution during 5 days and then exposed for 7 days to 1 µM Cs^+^. In order to alleviate detrimental effects of changing the solution composition, K^+^ concentration remained the same before and during exposure to Cs^+^. Some plants remained in the Cs^+^-free solution to analyse K^+^-content after growing in 10, 100 or 3000 µM K^+^-solution. Renewing of the solutions was performed every 2-3 days to avoid significant depletion in the medium due to uptake by plants.

For Rb^+^ uptake experiments, 30 days-old plants supplied with 10 µM K^+^-solution during 7 days were transferred in 20 mL of a K^+^-free solution containing 50 µM RbCl. After 7 h, roots were rinsed with Rb^+^-free solution to remove adsorbed Rb^+^.

### Cs^+^ toxicity assay

For evaluation of Cs^+^ effects on cotyledons development, surface-sterilized *Arabidopsis* seeds of Col-0, *hak5-3* (Qi et al., 2008), *kup9-1* and *kup9-3* were sown on 2 mL of nutritive solution containing 10 µM or 1000 µM K^+^. After 48h at 4 °C, 1 mL of nutritive solution contaminated with CsCl (final concentration ranging from 0 to 300 µM) was added and plants were allowed to grow during 7 days in growth chamber.

For evaluation of Cs^+^ effects on roots elongation, surface-sterilized *Arabidopsis* seeds of Col-0, *hak5-3* (Qi et al., 2008), *kup9-1* and *kup9-3* were sown and allowed to grow in MS½ agar plates vertically oriented during 4 days. Seedlings were then transferred under sterile conditions on different agar media containing K^+^ (10 or 1000 µM) and Cs^+^ (0, 10, 100, 300 or 500 µM). Primary root elongation was measured after 7 days on these plates, oriented vertically.

### Cs^+^ fluxes between seedlings and external solution

Protocol for Cs^+^ depletion experiments was adapted from Rb^+^ depletion experiments described in Rubio et al. (2008). Thirty days-old plants, supplied with 10 µM K^+^-solution during 7 days to stimulate Cs^+^ uptake, were transferred in 20 mL of a K^+^-free solution containing 60 µM CsCl. Cs^+^ depletion was followed taking up 100 µL of the solution at different time points during 24 h. Then, Cs^+^ contaminated plants were transferred in 20 mL of a 10 µM K^+^-nutrient solution containing no Cs^+^ after prior rinsing in this solution to remove adsorbed Cs^+^ bound to the roots. At different time points, 100 µL of the solution were taken to follow Cs^+^ released from plants to the external medium.

### Measure of Cs^+^, K^+^ and Rb^+^

Plants and aliquots of exposure solutions and of agar media were analysed for measuring and verifying K^+^, Cs^+^ and Rb^+^ concentrations. For plant samples, roots and shoot were separated, blotted on Benchkote paper and then oven dried (3-5 days at 50-60 °C). Aliquots (5 mL) of agar media and plants dried matters were mineralized in HNO_3_ 65% (5 or 10 mL for plants and agar media respectively) and H_2_O_2_ 30% (1.5 or 3 mL) at 100-150 °C on a sand bath. Mineralisates were evaporated to dryness and redissolved in HNO_3_ 2% v/v prior to analysis.

In all substrates, Rb^+^ and Cs^+^ were quantified by ICP-MS (Inductively Coupled Plasma-Mass Spectrometry, PQ Excell Thermo Electron with S-Option, detection limit 5 ng.L^−1^) and K^+^ content by ICP-AES (-Atomic Emission Spectrometry, OPTIMA 8300, Perkin Elmer, quantification limit 10 μg.L^−1^).

### Statistical analyses

ANOVA analyses were performed in the R environment (version 3.5.1) to evaluate effects of the different treatments on plant K^+^, Rb^+^ and Cs^+^ content and on root elongation separately (NS, Non-Significant and *, **, *** Significant at the α = 0.05, 0.01 and 0.001 level respectively). In tables, different letters in bold indicate significant differences between means (Tuckey post-hoc test, p-value < 0.05).

### Spatial transcription profiling of *KUP9*

The fragment of 2745 bp upstream of the start-codon of *KUP9* gene was PCR amplified (forward primer: CCAATGTAACGAGGGAAGAGACT, reverse primer: CAGGGGAATTTCGAGTTCTTTTGT) and then inserted into pCR-XL-TOPO^®^ vector. This first step allowed us to enhance subsequent amplification with *att*B primers (*att*B1 forward primer: *GGGGACAAGTTTGTACAAAAAAGCAGGCTAT*TGTAACGAGGGAAGAGACTTG, *att*B2 reverse primer *GGGGACCACTTTGTACAAGAAAGCTGGGTC*TTTTGTAACAAAAGAACTCGAAATTC) for *KUP9* promoter cloning using Gateway^®^ technology. Following the manufacturer’s instructions, *KUP9* promoter was introduced into the entry vector pDONR221™ and then cloned in pBGWFS7 plasmid containing a *GFP-GUS* fusion. The construct was introduced into *Agrobacterium tumefaciens* (C58C1) and transformed into Col-0 plants using the floral dip method (Clough & Bent, 1998). Staining of GUS activity was performed on T3 homozygous transgenic plants incubating tissues on a fixation solution (50 mM NaPO_4_ buffer-30 mM Na_2_HPO_4_ + 20 mM NaH_2_PO_4_-, 2 mM potassium ferricyanide, 2mM potassium ferrocyanide, 0,05% Triton X-100, 1 mg mL^−1^ X-Gluc, pH 7).

### Protein localisation analysis

KUP9 protein fusion with the GFP reporter was generated to localise the KUP9 transporter. The coding sequence of *KUP9* was PCR amplified (forward primer: ATGGCGGAAAGAGTCGAAGCATC, reverse primer: CTAAACATAAAAGACTTGTCCAACG) on Col-0 cDNA synthetized from 1 µg RNA with SuperScript™ III kit (Invitrogen). Sequencing of PCR products (GATC Biotech, Konstanz, Germany) reveals that the amplified fragments correspond to the splicing variant named At4g19960.2 in TAIR database. This coding sequence was inserted into PCR-XL-TOPO optimizing this way its further insertion into the entry vector pDONR P2r-P3 (*att*B2r forward primer: *GGGGACAGCTTTCTTGTACAAAGTGG****TC***ATGGCGGAAAGAGTCGAAG, *att*B3 reverse primer: *GGGGACAACTTTGTATAATAAAGTTG****C***CTAAACATAAAAGACTTGTCCAACG) for cloning using Gateway^®^ technology. One LR recombination was performed following the manufacturer’s instructions to combine simultaneously PDONR P2r-P3 containing the *KUP9* CDS, PDONR221 containing *GFP* CDS, PDONRP4-P1R containing either the *35S* promoter or the *KUP9* native promoter and the destination vector pB7m34GW. Transgenic *Arabidopsis* plants carrying the resulting pro*35S*:*GFP*-*KUP9* and pro*KUP9*:*GFP*-*KUP9* constructs were obtained using the floral dip method (Clough & Bent, 1998).

## RESULTS

### *KUP9* is preferentially expressed in roots and pollen and is affected by K^+^ supply

Reporter gene experiments were performed to determine spatial expression pattern of *KUP9* in plants. Transgenic *Arabidopsis* lines expressing a GFP-GUS fusion protein under the control of the native *KUP9* promoter (Pro*KUP*9:*GFP*-*GUS*) were generated. Following staining of different plant parts (**Fig. 2**), GUS coloration was observed mainly in 7 days-old seedlings roots and mature plants flowers. In leaves, GUS coloration was not detected in most cases or was restricted to small areas often close to the base of a trichome.

**Figure 2:**
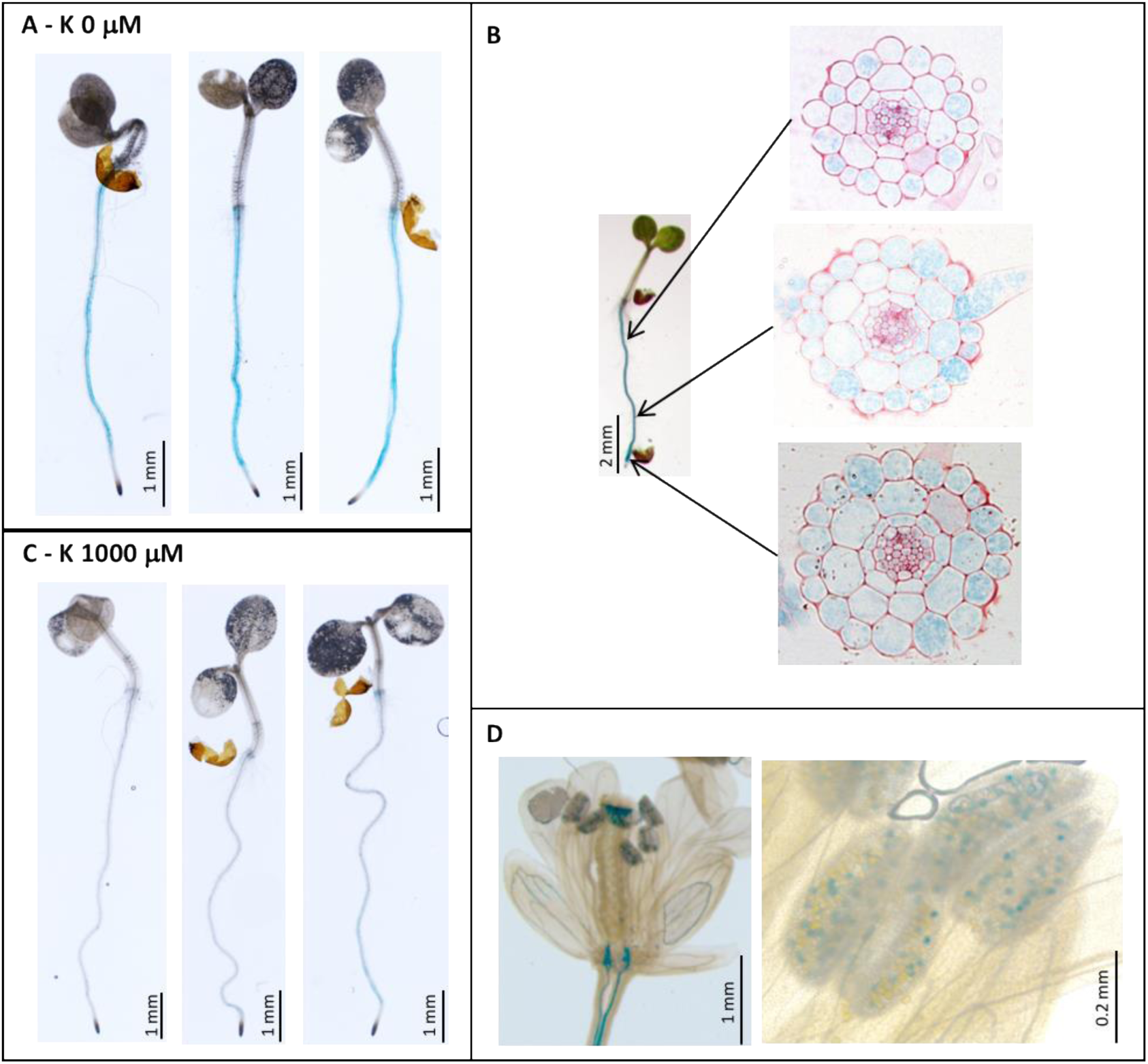
Spatial transcription pattern of *KUP9* determined in *ProKUP9:GFP-GUS* transgenic *A. thaliana* plants. GUS staining of (A) Roots of 7 days-old seedlings grown in nutritive solution containing no K^+^, (B) and cross section of a stained root, (C) Roots of 7 days-old seedlings grown in nutritive solution containing 1000 µM K^+^, (D) Flower and anthers of mature plants grown in sufficient K^+^-conditions.

In roots, GUS coloration was notably stronger in seedlings growing in nutritive solution containing low level of K^+^ (**Fig. 2.A and C)** suggesting that K^+^ supply affects *KUP9* transcription in 7 days-old *Arabidopsis* seedlings. This result was verified by quantitative PCR which indicated that *KUP9* expression is 1.4 fold higher (p-value 0.005) in Col-0 grown in low K^+^ supply compared to sufficient K^+^ condition (**Fig. S1**).

### KUP9 transporter localises to the cell membrane

Cellular localisation of the KUP9 transporter was achieved generating transgenic *A. thaliana* which expressed N-terminal GFP fusions with KUP9 transporter under the control of the 35S promoter. Confocal microscopy analysis of the GFP signal in transgenics lines revealed that KUP9 transporter is likely addressed to the plasma membrane (**Fig. 3**).

**Figure 3:**
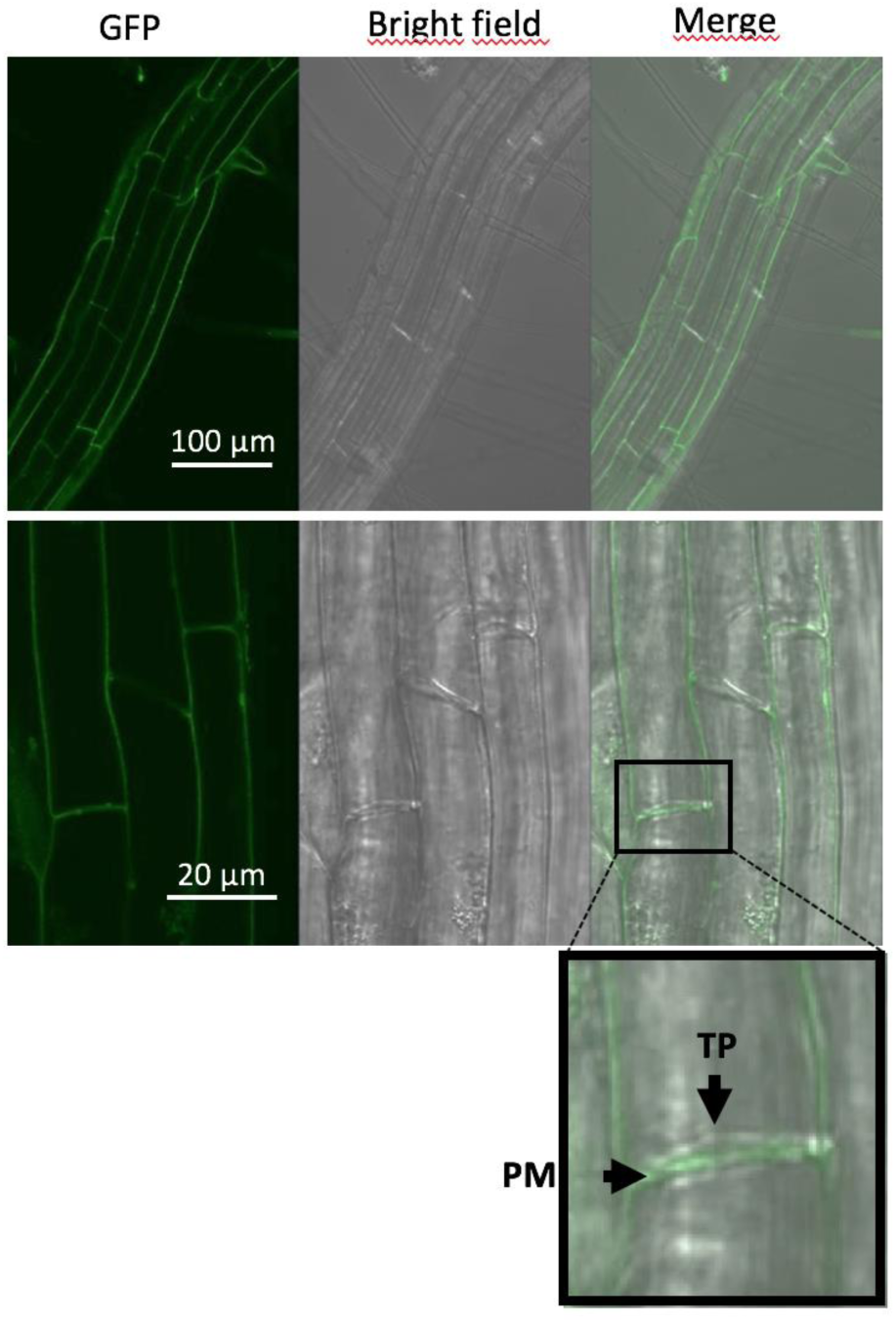
Localisation of KUP9 transporter observed in transgenic *A. thaliana* plants expressing the fusion *GFP-KUP9* under the control of the *35S* promoter. KUP9 transporter localises to the roots and is likely addressed to the plasma membrane. Arrows indicate the plasma membrane (PM) and tonoplast (TP).

As for *ProKUP*9:*GFP*-*GUS*, GFP signal was not detected in Pro*KUP*9:*GFP*-*KUP9* transgenic lines suggesting that the *KUP9* promoter activity was not sufficient.

### *kup9* mutants do not display defective K^+^ homeostasis

Different members of the *KUP/HAK/KT* family have been shown to be involved in K^+^ transport in *A. thaliana*. In the case of *AtKUP9*, there is no evidence *in planta* but it likely mediates K^+^ influx when expressed in an *Escherichia coli* mutant strain which lacks its three major K^+^ uptake systems (Kobayashi et al., 2010). To further understand its role in plant K^+^ transport, *kup9 Arabidopsis* mutant lines were compared to wild-type plants for their K^+^ status under three different levels of K^+^ supply (**Fig.4**).

**Figure 4:**
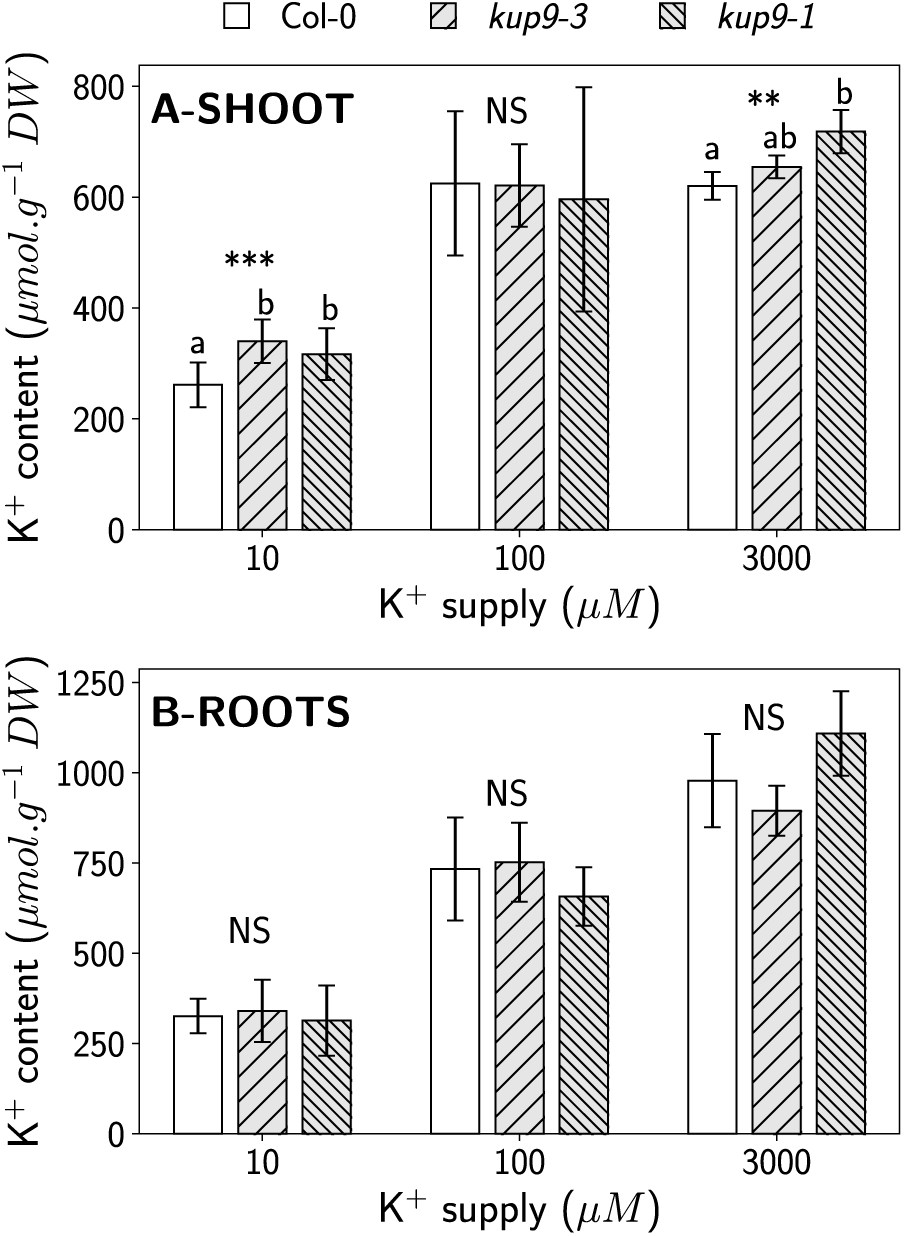
K^+^ content in 30 days-old plants supplied during 12 days with a nutrient solution containing 10, 100 or 3000 µM K^+^. Data for (A) Shoot and (B) Roots are means ± SD (n=11-12). ANOVA analyses were performed to compare K^+^ content between lines for each level of K^+^ supply (NS, Non-Significant; ** and *** Significant at the α = 0.01 and 0.001 level respectively). Different letters indicate significant differences between means (Tuckey post-hoc test, p-value < 0.05).

Under low (10 µM) and sufficient (3000 µM) K^+^ supply, shoot K^+^ content tended to be higher in *kup9* mutant lines compared to Col-0 whereas K^+^ content in roots were nearly the same for the three lines in all tested levels of K^+^ supply. In addition, there were no significant differences for Rb^+^ (used as a K^+^ tracer) influx in plants lacking *KUP9* compared to wild-type (**Table 1**). Taking together these results suggest that, in our conditions (30 days-old plants grown in nutrient solution containing 10, 100 or 3000 µM K^+^ during 7 days), disruption of *KUP9* does not significantly impaired K^+^ homeostasis (uptake and distribution).

**Table 1:** Rb^+^ accumulation in 30 days-old *A. thaliana* plants exposed during 7 h to 20 mL of a K^+^-free nutrient solution containing 50 µM RbCl. Before experiment, plants were supplied with a 10 µM K^+^ nutrient solution during 5 days to enhance subsequent Rb^+^ uptake. Data are means ± SD (n=5). Different letters indicate significant differences between means.

| Lines | Shoot Rb<sup>+</sup> content (μmol g<sup>-1</sup> DW) | Roots Rb<sup>+</sup> content (μmol g<sup>-1</sup> DW) |
| --- | --- | --- |
| Col-0 | 0.55 ± 0.20 (a) | 5.13 ± 0.95 (b) |
| *kup9-1* | 0.75 ± 0.30 (a) | 5.37 ± 1.37 (b) |
| *kup9-3* | 0.73 ± 0.30 (a) | 6.63 ± 0.95 (b) |

### *kup9* mutants are more sensitive to Cs^+^

Based on a previous work showing that a K^+^ transport-deficient *E. coli* mutant expressing *AtKUP9* was able to take up Cs^+^ (Kobayashi et al., 2010), we wondered whether *kup9 Arabidopsis* mutants differed from wild-type in their response to Cs^+^. Cotyledon development and primary root elongation of wild-type and of the two mutant lines disrupted in *AtKUP9* were compared on media containing low (10 µM) or high (1000 µM) level of K^+^ and a range of Cs^+^ concentrations (**Fig.5**). A mutant lacking *AtHAK5* previously screened in similar experiments (Qi et al., 2008) was used to validate our results.

**Figure 5:**
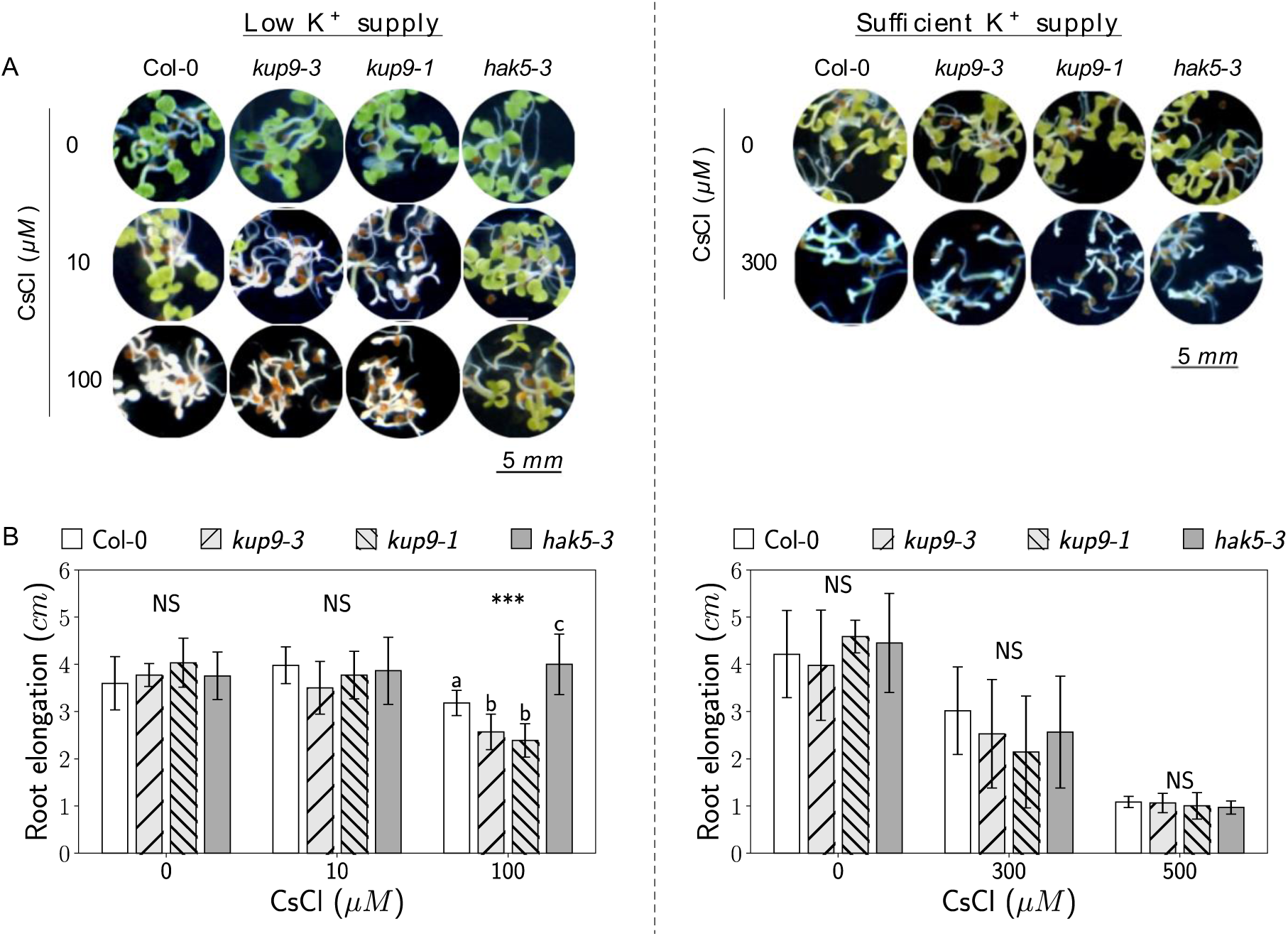
Sensitivity to Cs^+^ in *kup9* mutant lines. **(A)** 10 days-old seedlings grown in 8 mL of low K^+^ (10 µM) or sufficient K^+^ (1000 µM) nutritive solution and containing Cs^+^. **(B)** Primary root elongation of 11 days-old seedlings grown in agar plates in the presence of Cs^+^ and 10 µM K^+^ or 1000 µM K^+^. Plants were allowed to grow on MS½ agar plates during 4 days before transfer on Cs^+^. Data are means ± SD (n=10-16).

Cotyledons of seedlings grown in liquid media bleached in presence of toxic concentrations of Cs^+^ (**Fig.5.A**). Conversely to plants grown in media containing 1000 µM K^+^, the different lines displayed different sensitivities to Cs^+^ when grown in 10 µM K^+^. For *kup9* mutant lines supplied with low amount of K^+^, bleaching occurs at 10 µM of Cs^+^ whereas cotyledons of wild-type remain green. Increasing concentrations of Cs^+^ (100 µM), cotyledons of wild-type turned completely bleached whereas those of *hak5-3* remain partly green.

In vertical plates containing low amount of K^+^ (10 µM), primary root elongation of both *kup9* mutant lines was significantly lower than those of wild-type plants when Cs^+^ concentration reached 100 µM (**Fig.5.B**). Thus, plants lacking *KUP9* appeared more sensitive to Cs^+^ when grown under low K-supply. Conversely, *HAK5* disruption enhanced Cs^+^ tolerance under this same condition which is consistent with its role in Cs^+^ uptake. Discrepancies in Cs^+^ toxicity between the different lines were not visible in plants grown in higher amount of K^+^ (1000 µM).

### *kup9* mutants accumulate more Cs^+^

Toxicity of Cs^+^ in plants may originate from K^+^ starvation caused by the presence of Cs^+^ in the rhizosphere (Maathuis & Sanders, 1996) and is related to the Cs^+^:K^+^ concentration ratio at tissue level (Hampton et al., 2004). Since *KUP9* disruption does not alter negatively K^+^ status but does increase sensitivity to Cs^+^, we wondered whether it affects Cs^+^ accumulation *in planta*. Cs^+^ accumulation was compared in Col-0 wild-type and *kup9* mutant lines exposed to 1 µM CsCl during 7 days in hydroponics conditions (**Fig.6.A**). A *hak5* mutant line was also used to validate our experiments. In plants grown with sufficient K^+^ supply (3000 µM), disruption of *KUP9* and *HAK5* had no significant effect in Cs^+^ accumulation. When K^+^ supply was low (10 µM), Cs^+^ accumulation increased with different extent depending on the plant line. As expected under low K^+^ supply, plants lacking *HAK5* accumulated 50% less of Cs^+^ than the wild-type. In the same condition, disruption of *KUP9* had the opposite effect resulting in a higher accumulation of Cs^+^ than the wild-type (around 30% more). This is consistent with a recently published experiment performed in liquid media containing 500 µM K^+^ and 300 µM Cs^+^ showing that *kup9-1* seedlings accumulate more Cs^+^ than the wild-type Col-0 (Adams, Miyazaki & Shin, 2019).

**Figure 6:**
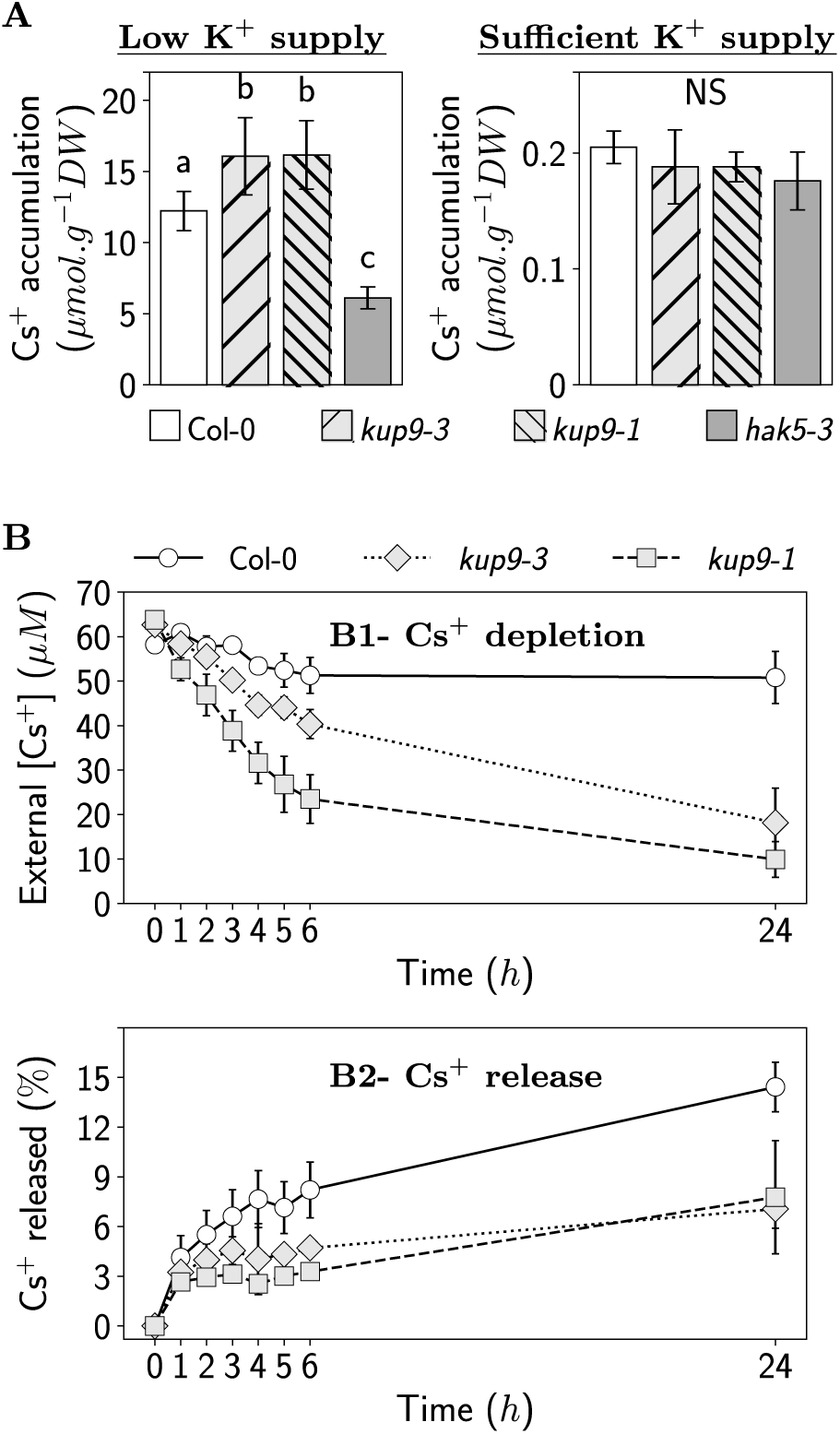
Accumulation of Cs^+^ in *kup9* mutant lines. **(A)** Cs^+^ accumulation in 30 days-old plants exposed during 7 days to a nutrient solution containing 1 µM Cs^+^ and 10 µM K^+^ or 3000 µM K^+^. Data are means ± SD (n=6-8). **(B)** Cs^+^ exchanges between 30 days-old plants and the external solution. External Cs^+^ concentrations were followed by taking up small samples of the solution. Data are means ± SD (n=5). (B1) Cs^+^ depletion due to uptake by plants in a K^+^-free solution. Before the experiment, plants were grown with low K^+^-supply (10 µM) during 7 days to improve Cs^+^ influx. (B2) Cs^+^ released from contaminated plants to a Cs^+^-free solution.

Examining separately shoot and roots Cs^+^ contents (**Table 2**), it appeared that Cs^+^ distribution remained globally unchanged in *kup9* mutant lines compared to wild-type. Interestingly, when K^+^-supply was low (10 µM), *hak5-3* mutants displayed a divergent pattern of Cs^+^ distribution with a greater part of Cs^+^ allocated to shoot. An impairment of the systems involved in Cs^+^ translocation or redistribution in plants lacking *HAK5* seems unlikely since HAK5 transporter has been demonstrated to be involved in roots Cs^+^ influx (Qi et al., 2008). Therefore, this result could be due to the overall decrease of Cs^+^ accumulation described above in *hak5-3* mutants.

**Table 2:**
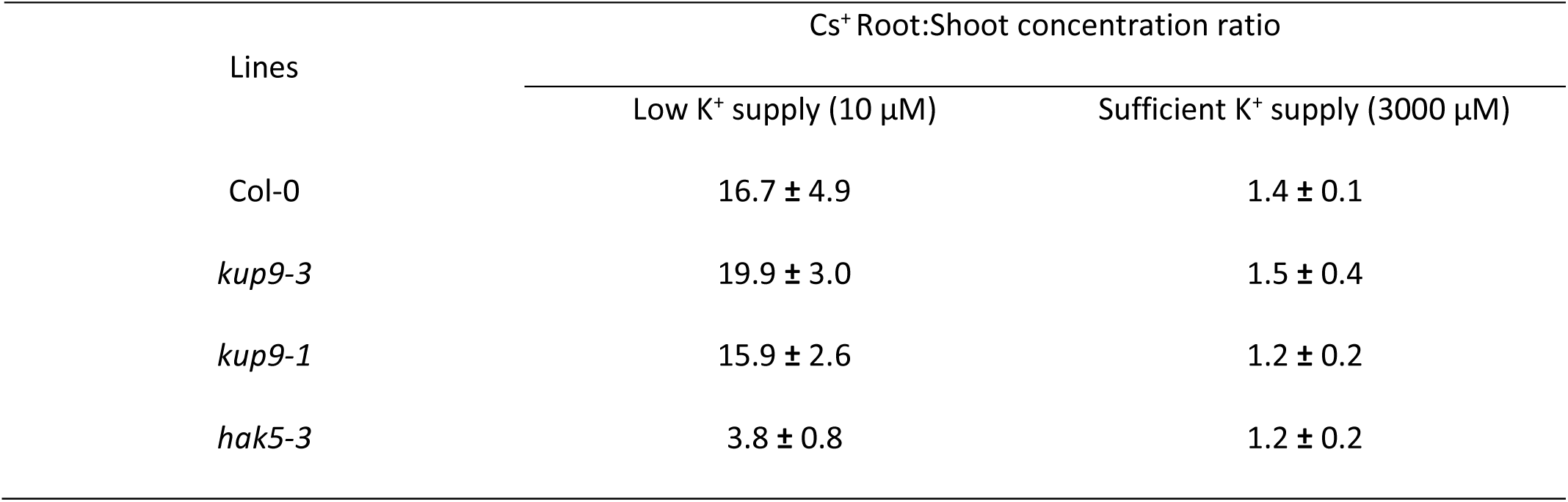
Cs^+^ roots:shoot concentration ratio in plants exposed during 7 days to a nutrient solution containing 1 µM CsCl and 10 µM K^+^ or 3000 µM K^+^. Data are means ± SD (n=6-8).

| Lines | Cs<sup>+</sup> Root:Shoot concentration ratio |  |
| --- | --- | --- |
|  | Low K<sup>+</sup> supply (10 μM) | Sufficient K<sup>+</sup> supply (3000 μM) |
| Col-0 | 16.7 ± 4.9 | 1.4 ± 0.1 |
| *kup9-3* | 19.9 ± 3.0 | 1.5 ± 0.4 |
| *kup9-1* | 15.9 ± 2.6 | 1.2 ± 0.2 |
| *hak5-3* | 3.8 ± 0.8 | 1.2 ± 0.2 |

To further investigate mechanisms leading to an increase of Cs^+^ accumulation in plants lacking *KUP9*, fluxes of Cs^+^ between roots and the external medium were measured in wild-type and in *kup9* mutant lines (**Fig.6**). First, K^+^-starved plants were exposed to a K^+^-free solution containing Cs^+^. In 24 h, amount of Cs^+^ in this solution decreased by 74 to 85% in *kup9* mutant lines whereas only 30% was taken up by wild-type plants. Divergences between lines appeared within the first three hours but are not clear during the first hour suggesting that effects of *KUP9* disruption on Cs^+^ transport is not immediate. Contaminated plants were then transferred in a Cs^+^-free solution in which Cs^+^ release was followed. Significant amount of Cs^+^ was detected in the external medium from the first hour suggesting that Cs^+^ is quickly released by plants. After 24 h, the initially Cs^+^-free solution contained 2.71 µM (± 0.57), 4.07 µM (± 0.39) and 3.47 µM (± 1.15) of Cs^+^ for Col-0, *kup9-3* and *kup9-1* respectively. Interestingly, it was noticed that *kup9* lines released less Cs^+^ than wild-type in proportion when amount of Cs^+^ released in the solution was normalized by amount of Cs^+^ accumulated in plants at the beginning of experiment.

## DISCUSSION

### KUP9 prevents Cs^+^ accumulation in plants

Cs^+^ accumulation in plants depends on the level of K^+^-supply and several K^+^ transporters have been shown to mediate opportunistic Cs^+^ fluxes (**Fig. 7**). Role of HAK5 (Qi et al., 2008) and NSCC (Hampton et al., 2005; White & Broadley, 2000) in root Cs^+^ uptake and regulation of Cs^+^ content in xylem by ZIFL2 (Remy et al., 2015) have partly deciphered the molecular entities involved in Cs^+^ transport. In contrast, identities of carriers involved in Cs^+^ transport from cytosol to intracellular compartments as well as mechanisms underlying Cs^+^ release from root cells to the external medium remain unknown. In the present study, analysis of *kup9* mutant lines provide several points of evidence involving KUP9 transporter in the limitation of Cs^+^ accumulation under low K^+^-supply condition in *A.thaliana*.

**Figure 7:**
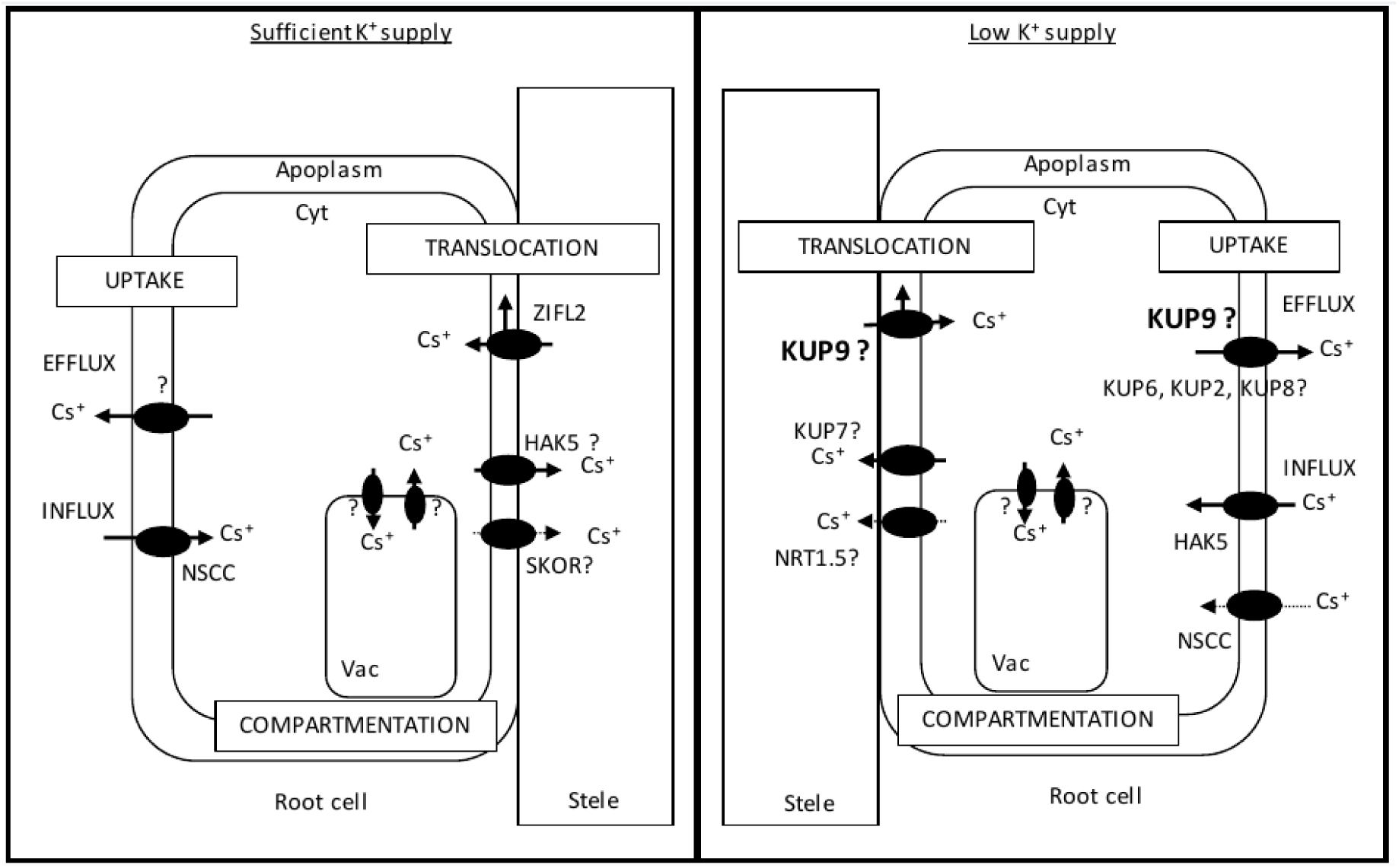
Model for Cs^+^ transport in roots of *A. thaliana* depending on the level of K^+^ supply. INFLUX. As for other cations, Cs^+^ taken up by plants from the soil solution is transported across the root mainly through the symplastic pathway. Non-selective Cation Channels (NSCC, Hampton et al., 2005; White & Broadley, 2000) and the high-affinity K^+^ transporter HAK5 (Qi et al, 2008) are thought to mediate the major part of Cs^+^ influx in sufficient (mM range) and low (μM range) K^+^ conditions respectively. TRANSLOCATION. Cs^+^ is highly mobile within plants. From the roots, Cs^+^ is loaded into the xylem and distributed towards the aerial parts through yet unidentified transport systems. In this way, further investigations are needed to determine roles of transport systems involved in K^+^ translocation such as the stellar K^+^ outward-rectifying channel SKOR (Drechsler et al., 2015; Gaymard et al., 1998), the nitrate transporter1/peptide transporter NRT1.5 (Drechsler et al., 2015; Li et al., 2017), the K^+^ transporters KUP7 (Han et al., 2016) and HAK5 (Nieves-Cordones et al., 2019). The Major Facilitator Superfamily transporter ZIFL2 has been proposed to prevent Cs^+^ xylem loading mediating Cs^+^ release into the apoplasm of endodermis and pericycle, under Cs^+^ and K^+^ excess (Remy et al., 2015). COMPARTMENTATION. Regarding its intracellular distribution, analyses of contaminated plant cells revealed that Cs^+^ is not limited to the cytosol but is also found in vacuole and probably chloroplasts (Akamatsu et al., 2014; Le Lay et al., 2006). Identities of transport systems involved in these mechanisms are unknown. EFFLUX. Our results suggest that KUP9 prevents plant Cs^+^ accumulation promoting its release from root cells. This is supported by a previous analysis of the triple mutant *kup268* which suggests that these KUP might be involved in root K^+^ efflux. Similarly to the ZIFL2 transporter in sufficient K^+^ condition, KUP9 could also participate to Cs^+^ release into the apoplasm before release into the external medium in low K^+^ condition. Cyt: cytosol, Vac: Vacuole.

First, *kup9* mutant lines display higher sensitivity to Cs^+^ which is reflected by cotyledons bleaching and reduced root elongation for lower Cs^+^ concentrations compared to wild-type. When used in the millimolar range, Cs^+^ induces K^+^ decrease in shoots of intoxicated plants (Hampton et al., 2004). In our test, micromolar Cs^+^ concentrations were sufficient to point out the higher sensitivity to Cs^+^ in *kup9* mutant lines and we did not measure significant decrease of K^+^ status in shoot of Cs^+^-intoxicated plants in these conditions (**Fig.S2**). Higher toxicity of Cs^+^ in *kup9* mutant lines is therefore likely linked to the toxicity of Cs^+^ itself rather than to a concomitant alteration of K^+^ status in shoot. This is supported by the second point showing the role of KUP9 in Cs^+^ transport, which is the higher accumulation of Cs^+^ in *kup9* mutant lines. Conversely to results showing that *AtKUP9* mediates Cs^+^ uptake when expressed in *E. coli* mutants (Kobayashi et al., 2010), our results demonstrate therefore that *KUP9* is not directly involved in Cs^+^ influx in roots of *A. thaliana* but participate to the limitation of Cs^+^ accumulation in plants. Such discrepancies between heterologous and *in planta* analyses have been discussed elsewhere and imputed for instance to artificial interactions with heterologous structures or to the lack of regulatory proteins from the native organism (Dreyer et al., 1999).

Level of K^+^-supply has a major effect on Cs^+^ accumulation in plants. This is linked to several factors such as the competition between the two cations for the same transport systems and the control of transporter expression by K^+^-supply. In our study, *kup9* mutants displayed enhanced Cs^+^ accumulation only when supplied with low level of K^+^. We suggest that, at sufficient K^+^-supply (mM range), relative lower transcription of *KUP9* in wild-type plants and higher external Cs^+^ dilution may blur discrepancies between *kup9* and wild-type Cs^+^ accumulation.

In the present study, we have shown that Cs^+^ release from roots to external solution is proportionally two times lower in plants lacking *KUP9* compared to wild-type. Taking this result together with the localisation of KUP9 transporter in root cells of *pro35S:GFP-KUP9* transgenic lines, we propose that KUP9 participate to the efflux of cations from roots (**Fig. 7**). This is supported by similar results suggesting that the KUP6 sub-family, i.e. KUP6, KUP8 and KUP2 (**Fig. 8**), may participate with the Shaker channel GORK in the efflux of K^+^ from roots during K^+^ starvation (Osakabe et al., 2013).

**Figure 8:**
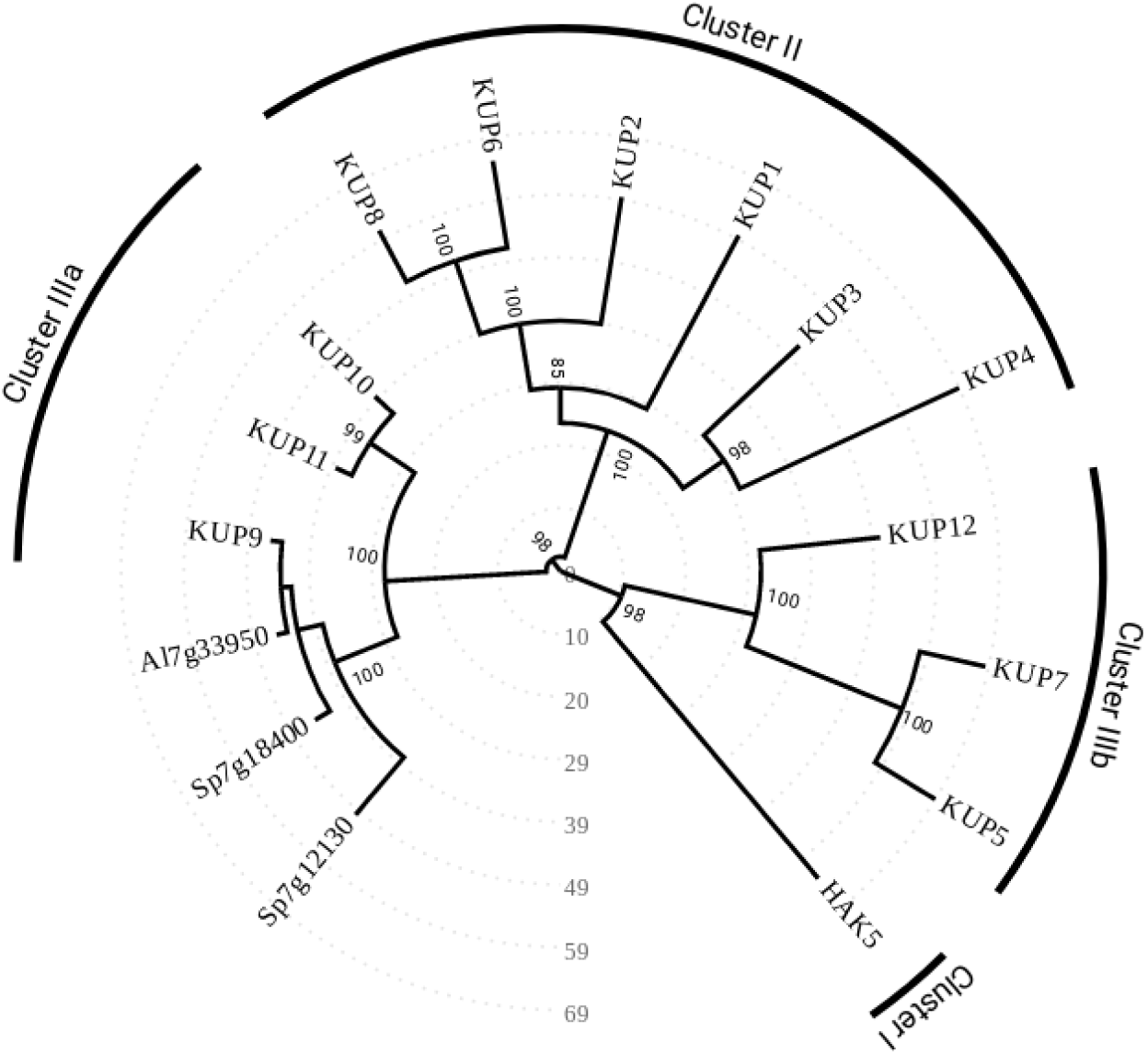
Phylogenetic organization of the KUP/HAK/KT transporters family in *Arabidopsis thaliana*. The phylogenetic tree was generated from polypeptide sequences with the PhyML software (v3.0) in the online platform Phylogeny.fr (« One Click » mode with Muscle for alignment and Gblock for alignment curation, http://www.phylogeny.fr/simple_phylogeny.cgi). The unrooted tree was drawn with the online software PONYTREE (website under construction). Bootstrap values (in percentage) are indicated at the corresponding nodes and the scale represents the number of changes per 100 amino acid. Polypeptide sequences of AtKUP9 homologs (Al7g33950, Sp7g18400 and Sp7g12130) and AtKUP/HAK/KT were from thellungiella.org (http://thellungiella.org), Phytozome (https://phytozome.jgi.doe.gov/pz/portal.html) and TAIR database (https://www.arabidopsis.org/).

### Primary plant role of KUP9

Cs^+^ has no relevant functions in plants suggesting that transport of Cs^+^ through KUP9 transporter is more probably a non-specific process. In *A. thaliana*, KUP9 belongs to the KUP/HAK/KT family organized in 4 clades (**Fig. 8**) and the others KUP transporters have been related to various biological processes such as roots K^+^ acquisition (HAK5, Gierth et al., 2005; Rubio et al., 2000) and translocation (KUP7, Han, Wu, Wu & Wang, 2016) or growth and development (KUP4, Rigas et al., 2001 and KUP2, Elumalai, Nagpal & Reed, 2002, KUP6 and KUP8, Osakabe et al., 2013). In *E. coli* mutants defective for constitutive K^+^ transport systems, heterologous expression of *AtKUP9* is able to mediate K^+^ uptake (Kobayashi et al., 2010) but there is no published *in planta* evidence involving KUP9 in *A. thaliana* K^+^ homeostasis to our knowledge. In others plant species, such as the extremophytes *Schrenkiella parvula, A. lyrata* and *A. arenosa*, it is thought that higher expression strength and single-nucleotide polymorphism affecting *KUP9* homologs might be related to an adjustment in K^+^ transport supporting plants adaptation to soils containing challenging ions concentrations (Arnold et al., 2016; Oh et al., 2014; Turner, Bourne, Von Wettberg, Hu & Nuzhdin, 2010). For *A. thaliana* tested under experimental conditions described here, however, there were no significant effects of *KUP9* disruption in K^+^ homeostasis. At the contrary of Cs^+^, K^+^ is involved in many biological processes and K^+^ homeostasis is tightly regulated. Therefore, compensatory mechanisms through redundant functions of K^+^ carriers could be induced in the tested *kup9* mutant lines blurring potential discrepancies with wild-type plants. Based on the phylogenetic relationship between KUP9, KUP10 and KUP11 (**Fig. 8**) it could be interesting to investigate the triple mutant as it has been done for KUP2, KUP6 and KUP8 pointing out their role in K^+^ efflux (Osakabe et al., 2013).

Primary plant role of KUP9 could also be significant in other plant stages not tested in this study. Activity of GUS reporter in transgenic plants carrying the Pro*KUP*9:*GFP*-*GUS* construct reveals expression of *KUP9* in certain pollen grains (**Fig. 2D**). This is consistent with previous transcriptome analyses showing the late pollen-expression pattern of *KUP9* which differs from the other *KUP* on this point (Bock et al., 2006). Functions of potassium transporters are crucial for pollen tube growth (Mouline et al., 2002) and it could be interesting to further investigate the role of *KUP9* in pollen development.

## CONCLUSION

The use of plants for remediation of radiocesium-contaminated soils as well as development of “safe crops” receive considerable interest since few decades (White et al., 2003; Zhu & Shaw, 2000). However, the fact that Cs^+^ enters plants through K^+^ transport system is a major constraint for phytoremediation strategies. Controlling expression of major potassium transporters, such as HAK5, to increase/decrease plant Cs^+^ acquisition without disturbing K^+^ nutrition may imply to modulate their specificity (Alemán et al., 2014). By contrast, the two *kup9* mutant lines tested in the present study display higher Cs^+^ accumulation without significant alteration of K^+^ status when supplied with low level of K^+^. We proposed that KUP9 may prevent Cs^+^ accumulation releasing it from root cells and that the potential role of KUP9 in K^+^ homeostasis appears to be minor in experimental conditions used here. Manipulating expression of such minor K^+^ transporters, whose disruption does not alter plant K^+^ acquisition but does enhance substantially Cs^+^ accumulation, may offer a valuable alternative for phytoremediation strategies.

## AKNOWLEDGMENTS

This work was cofunded by IRSN and CEA and has benefited from funds of the French government, managed by the Agence Nationale de la Recherche, originating from the funding program “Investissement d’Avenir” under the reference ANR-11-RSNR-0005.

## SUPPLEMENTAL FIGURES

**Figure S1:**
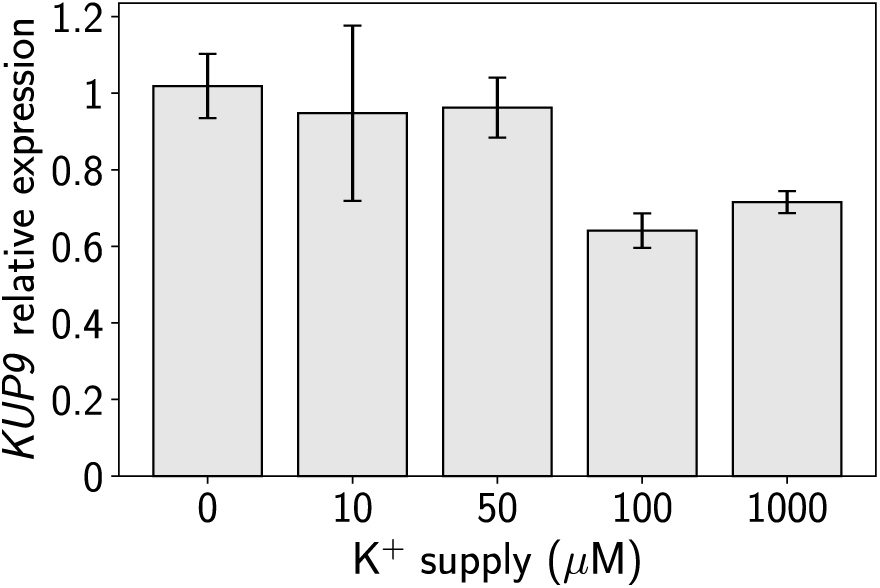
*KUP9* relative expression measured in RT-qPCR. Total RNA of roots from Col-0 supplied with different amount of K^+^ (0, 10, 50, 100 or 1000 µM) was extracted using the RNeasy™ kit (QIAgen) according to the manufacturer’s instructions. Reverse transcription of mRNA was performed over 1 µg of total RNA using SuperScript™ Vilo™ kit (Invitrogen) with oligo(dT)_20_ primers. The synthesized cDNAs were analysed by quantitative real-time PCR using SYBR^®^ Green I Master mix on a LightCycler^®^ 480 (Roche). PCR amplifications of a *KUP9* fragment (forward primer: AGAGGAGGAGGAGACGGATGAG, reverse primer: GCCCTACAAATCTTAGCAAG) were performed at 95°C for 10 sec (45 cycles) and 60°C for 10 sec. Relative quantitative results were calculated after normalization to ROC3. Data are mean +/-SD of a pool of three plants analysed three times.

**Figure S2:**
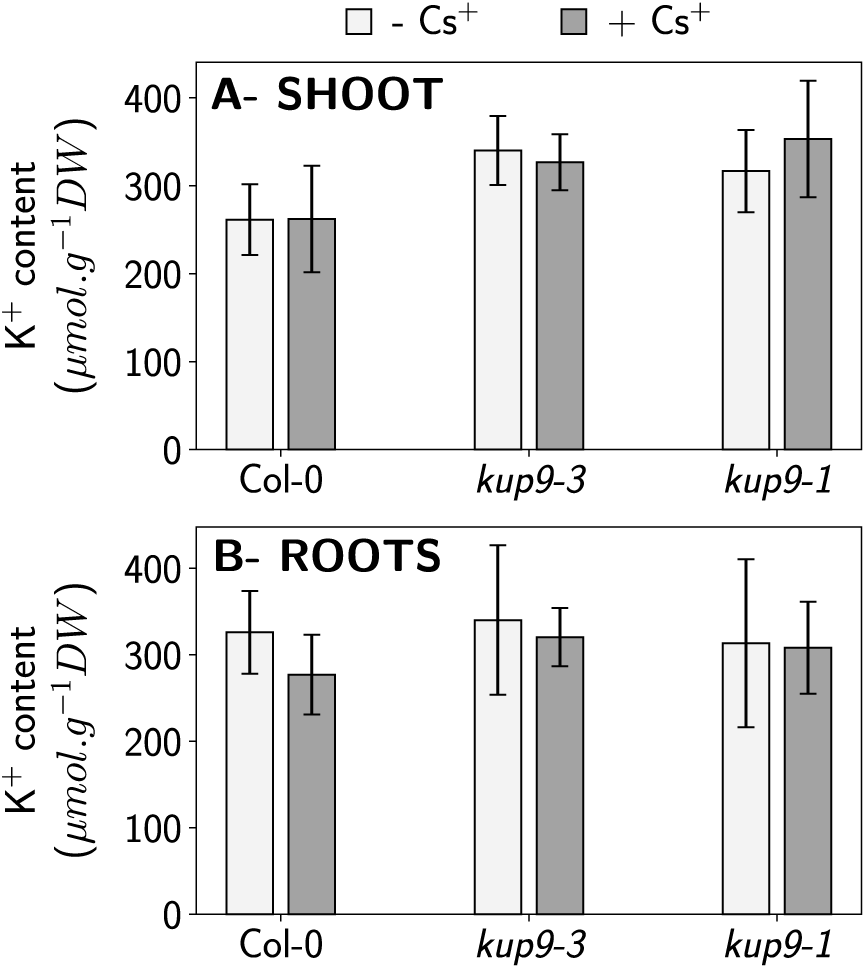
K^+^ content in Cs^+^-exposed plants. K^+^ content was measured in 30 days-old plants supplied during 12 days with a nutrient solution containing 10 µM K^+^ and exposed during 7 days to 1 µM Cs^+^ (+ Cs^+^) or not exposed (-Cs^+^). Data for (A) Shoot and (B) Roots are means ± SD (n=11-12). ANOVA analyses performed to compare K^+^ content between lines for each level of K^+^ supply revealed no significant differences.

## REFERENCES

Adams, E., Miyazaki, T., & Shin, R. (2019). Contribution of KUPs to potassium and cesium accumulation appears complementary in *Arabidopsis*. Plant Signaling & Behavior, 14: 1554468

Akamatsu, M., Komatsu, H., Mori, T., Adams, E., Shin, R., Sakai, H., Abe, M., Hill, J.P., & Ariga, K. (2014). Intracellular Imaging of Cesium Distribution in *Arabidopsis* Using Cesium Green. ACS applied materials & interfaces, 6, 8208–8211

Alemán, F., Nieves-Cordones, M., Martínez, V., & Rubio, F. (2011). Root K^+^ acquisition in plants: the *Arabidopsis thaliana* model. Plant and Cell Physiology, 52, 1603–1612

Alemán, F., Caballero, F., Ródenas, R., Rivero, R.M., Martínez, V., & Rubio, F. (2014). The F130S point mutation in the Arabidopsis high-affinity K^+^ transporter AtHAK5 increases K^+^ over Na^+^ and Cs^+^ selectivity and confers Na^+^ and Cs^+^ tolerance to yeast under heterologous expression. Frontiers in Plant Science, 5, 430

Arnold, B.J., Lahner, B., DaCosta, J.M., Weisman, C.M., Hollister, J.D., Salt, D.E., Bomblies, K., & Yant, L. (2016). Borrowed alleles and convergence in serpentine adaptation. Proceedings of the National Academy of Sciences, 113, 8320–8325

Bange, G., & Overstreet, R. (1960). Some observations on absorption of cesium by excised barley roots. Plant Physiology, 35, 605

Bertl, A., Reid, J.D., Sentenac, H., & Slayman, C.L. (1997). Functional comparison of plant inward-rectifier channels expressed in yeast. Journal of Experimental Botany, 48, 405–413

Bock, K.W., Honys, D., Ward, J.M., Padmanaban, S., Nawrocki, E.P., Hirschi, K.D., Twell, D., & Sze, H. (2006). Integrating membrane transport with male gametophyte development and function through transcriptomics. Plant Physiology, 140, 1151–1168

Bossemeyer, D., Schlösser, A., & Bakker, E.P. (1989). Specific cesium transport via the *Escherichia coli* Kup (TrkD) K^+^ uptake system. Journal of bacteriology, 171, 2219–2221

Bowen, H.J.M. (1979) Environmental chemistry of the elements. Academic Press, London.

Broadley, M.R., Escobar-Gutiérrez, A.J., Bowen, H.C., Willey, N.J., & White, P.J. (2001). Influx and accumulation of Cs^+^ by the *akt1* mutant of *Arabidopsis thaliana* (L.) Heynh. lacking a dominant K^+^ transport system. Journal of Experimental Botany, 52, 839–844

Clough, S.J., & Bent, A.F. (1998). Floral dip: A simplified method for *Agrobacterium*-mediated transformation of *Arabidopsis thaliana*. The Plant Journal, 16, 735–743.

Collander, R. (1941). Selective absorption of cations by higher plants. Plant Physiology, 16, 691

Dreyer, I., Horeau, C., Lemaillet, G., Zimmermann, S., Bush, D.R., Rodríguez-Navarro, A., Schachtman, D.P., Spalding, E.P., Sentenac, H., & Gaber, R.F. (1999). Identification and characterization of plant transporters using heterologous expression systems. Journal of Experimental Botany, 50, 1073–1087

Drechsler, N., Zheng, Y., Bohner, A., Nobmann, B., von Wirén, N., Kunze, R., & Rausch, C. (2015). Nitrate-dependent control of shoot K homeostasis by NPF7.3/NRT1.5 and SKOR in Arabidopsis. Plant Physiology, 169, 2832–2847.

Elumalai, R.P., Nagpal, P., & Reed, J.W. (2002). A mutation in the *Arabidopsis KT2/KUP2* potassium transporter gene affects shoot cell expansion. The Plant Cell Online, 14, 119–131

Epstein, E., & Hagen, C. (1952). A kinetic study of the absorption of alkali cations by barley roots. Plant Physiology, 27, 457

Gaymard, F., Pilot, G., Lacombe, B., Bouchez, D., Bruneau, D., Boucherez, J., Michaux-Ferrière, N., Thibaud, J.B., Sentenac, H. (1998). Identification and disruption of a plant shaker-like outward channel involved in K+ release into the xylem sap. Cell, 94, 647–655.

Genies, L., Orjollet, D., Carasco, L., Camilleri, V., Frelon, S., Vavasseur, A., Leonhardt, N., & Henner, P. (2017). Uptake and translocation of cesium by *Arabidopsis thaliana* in hydroponics conditions: Links between kinetics and molecular mechanisms. Environmental and Experimental Botany, 138, 164–172

Gierth, M., Mäser, P., & Schroeder, J.I. (2005). The potassium transporter AtHAK5 functions in K^+^ deprivation-induced high-affinity K^+^ uptake and AKT1 K^+^ channel contribution to K^+^ uptake kinetics in *Arabidopsis* roots. Plant Physiology, 137, 1105–1114

Greenwood, N.N., & Earnshaw, A. (1984). Chemistry of the Elements. Pergamon Press, Oxford.

Hamada, N. & Ogino, H. (2012). Food safety regulations: what we learned from the Fukushima nuclear accident. Journal of environmental radioactivity, 111, 83–99

Hampton, C.R., Broadley, M.R., & White P.J. (2005). Short review: the mechanisms of radiocaesium uptake by *Arabidopsis* roots. Nukleonika, 50, 3–8

Hampton, C.R., Bowen, H.C., Broadley, M.R., Hammond, J.P., Mead, A., Payne, K. A., Pritchard, J., & White, P.J. (2004). Cesium toxicity in Arabidopsis. Plant Physiology, 136, 3824–3837

Han, M., Wu, W., Wu, W.-H., & Wang, Y. (2016). Potassium Transporter KUP7 Is Involved in K^+^ Acquisition and Translocation in *Arabidopsis* Root under K^+^-Limited Conditions. Molecular plant, 9, 437–446

Hirsch, R.E., Lewis, B.D., Spalding, E.P., & Sussman, M.R. (1998). A role for the AKT1 potassium channel in plant nutrition. Science, 280, 918–921

Kobayashi, D., Uozumi, N., Hisamatsu, S., & Yamagami, M. (2010). AtKUP/HAK/KT9, a K^+^ transporter from *Arabidopsis thaliana*, mediates Cs^+^ uptake in *Escherichia coli*. Bioscience, biotechnology, and biochemistry, 74, 203–205

Kondo, M., Makino, T., Eguchi, T., Goto, A., Nakano, H., Takai, T., Arai-Sanoh, Y., & Kimura, T. (2015). Comparative analysis of the relationship between Cs and K in soil and plant parts toward control of Cs accumulation in rice. Soil Science and Plant Nutrition, 61, 144–151

Le Lay, P., Isaure, M.-P., Sarry, J.-E., Kuhn, L., Fayard, B., Le Bail, J.-L., Bastien, O., Garin, J., Roby, C., & Bourguignon, J. (2006). Metabolomic, proteomic and biophysical analyses of *Arabidopsis thaliana* cells exposed to a caesium stress. Influence of potassium supply. Biochimie, 88, 1533–1547

Li, H., Yu, M., Du, X. Q., Wang, Z. F., Wu, W. H., Quintero, F. J., Jin, X. H., Li, H.D. & Wang, Y. (2017). NRT1.5/NPF7.3 functions as a proton-coupled H^+^/K^+^ antiporter for K^+^ loading into the xylem in *Arabidopsis*. Plant Cell%, 29, 2016–2026.

Maathuis, F.J., & Sanders, D. (1996). Characterization of *csi52*, a Cs^+^ resistant mutant of *Arabidopsis thaliana* altered in K^+^ transport. The Plant Journal, 10, 579–589

Mäser, P., Thomine, S., Schroeder, J.I., Ward, J.M., Hirschi, K., Sze, H., Talke, I.N., Amtmann, A., Maathuis, F.J., & Sanders, D. (2001). Phylogenetic relationships within cation transporter families of *Arabidopsis*. Plant Physiology, 126, 1646–1667

Middleton, L.J., Handley, R., & Overstreet, R. (1960). Relative uptake and translocation of potassium and cesium in barley. Plant Physiology, 35, 913

Mouline, K., Véry, A.A., Gaymard, F., Boucherez, J., Pilot, G., Devic, M., Bouchez, D., Thibaud, J.B., & Sentenac, H. (2002). Pollen tube development and competitive ability are impaired by disruption of a Shaker K^+^ channel in *Arabidopsis*. Genes & Development%, 16, 339–350

Murashige, T., & Skoog, F. (1962). A revised medium for rapid growth and bio assays with tobacco tissue cultures. Physiologia Plantarum, 15, 473–497

Nieves-Cordones, M., Lara, A., Ródenas, R., Amo, J., Rivero, R.M., Martínez, V., & Rubio, F. (2019). Modulation of K^+^ translocation by AKT1 and AtHAK5 in *Arabidopsis* plants. Plant, Cell & Environment, 42, 2357–2371.

Oh, D.-H., Hong, H., Lee, S.Y., Yun, D.-J., Bohnert, H.J., & Dassanayake, M. (2014). Genome structures and transcriptomes signify niche adaptation for the multiple-ion-tolerant extremophyte *Schrenkiella parvula*. Plant physiology, 164, 2123–2138

Osakabe, Y., Arinaga, N., Umezawa, T., Katsura, S., Nagamachi, K., Tanaka, H., Ohiraki, H., Yamada, K., Seo, S.U., Abo, M., Yoshimura, E., Shinozaki, K., & Yamaguchi-Shinozaki, K. (2013) Osmotic stress responses and plant growth controlled by potassium transporters in *Arabidopsis*. Plant Cell, 25, 609–624

Qi, Z., Hampton, C.R., Shin, R., Barkla, B.J., White, P.J., & Schachtman, D.P. (2008). The high affinity K^+^ transporter AtHAK5 plays a physiological role *in planta* at very low K^+^ concentrations and provides a caesium uptake pathway in *Arabidopsis*. Journal of Experimental Botany, 59, 595–607

Remy, E., Cabrito, T.R., Batista, R.A., Teixeira, M.C., Sa-Correia, I., & Duque, P. (2015). The Major Facilitator Superfamily Transporter ZIFL2 Modulates Cesium and Potassium Homeostasis in *Arabidopsis*. Plant and Cell Physiology, 56, 148–162

Rigas, S., Debrosses, G., Haralampidis, K., Vicente-Agullo, F., Feldmann, K.A., Grabov, A., Dolan, L., & Hatzopoulos, P. (2001). *TRH1* encodes a potassium transporter required for tip growth in *Arabidopsis* root hairs. Plant Cell, 13, 139–151

Rubio, F., Guillermo, E., & Rodríguez-Navarro, A. (2000). Cloning of *Arabidopsis* and barley cDNAs encoding HAK potassium transporters in root and shoot cells. Physiologia Plantarum, 109, 34–43

Rubio, F., Nieves-Cordones, M., Alemán, F., & Martínez, V. (2008). Relative contribution of AtHAK5 and AtAKT1 to K+ uptake in the high-affinity range of concentrations. Physiologia Plantarum, 134, 598–608.

Sacchi, G.A., Espen, L., Nocito, F., & Cocucci, M. (1997). Cs^+^ uptake in subapical maize root segments: Mechanism and effects on H^+^ release, transmembrane electric potential and cell pH. Plant and Cell Physiology,38, 282–289

Shaw, G., & Bell, J. (1989). The kinetics of caesium absorption by roots of winter wheat and the possible consequences for the derivation of soil-to-plant transfer factors for radiocaesium. Journal of Environmental Radioactivity, 10, 213–231

Smolders, E., Kiebooms, L., Buysse, J., & Merckx, R. (1996). 137Cs uptake in spring wheat (*Triticum aestivum* L. cv. Tonic) at varying K supply. Plant and soil, 181, 211–220

Steinhauser, G., Brandl, A., & Johnson, T.E. (2014). Comparison of the Chernobyl and Fukushima nuclear accidents: A review of the environmental impacts. Science of the total environment, 470, 800–817

Tenorio-Berrío, R., Pérez-Alonso, M.-M., Vicente-Carbajosa, J., Martín-Torres, L., Dreyer, I. & Pollmann, S. (2018). Identification of two auxin-regulated potassium transporters involved in seed maturation. International Journal of Molecular Sciences, 19, 2132.

Turner, T.L., Bourne, E.C., Von Wettberg, E.J., Hu, T.T., & Nuzhdin, S.V. (2010). Population resequencing reveals local adaptation of *Arabidopsis lyrata* to serpentine soils. Nature genetics, 42, 260–263

White, P.J., & Broadley, M.R. (2000). Tansley Review No. 113. New Phytologist, 147, 241–256

White, P.J., Swarup, K., Escobar-Gutiérrez, A.J., Bowen, H.C., Willey, N.J., & Broadley, M.R. (2003). Selecting plants to minimise radiocaesium in the food chain. Plant and soil, 249, 177–186

Zhu, Y.G., & Shaw, G. (2000). Soil contamination with radionuclides and potential remediation. Chemospher, 41, 121–128

